# Multi-scale modelling of shear stress on the syncytiotrophoblast: Could maternal blood flow impact placental function across gestation?

**DOI:** 10.1101/2022.06.12.495853

**Authors:** Tet Chuan Lee, Ali Moulvi, Joanna L. James, Alys R. Clark

## Abstract

The surface of the placenta is lined by a single multinucleated cell, the syncytiotrophoblast, which forms a functional barrier between maternal and fetal blood in pregnancy. The placenta plays a critical role in healthy fetal development and over the course of pregnancy forms a complex branching tree-like structure which bathes in maternal blood and serves a vital exchange function. It has been suggested that the structure of the placenta may evolve, in part, under the influence of the shear stress exerted by maternal blood flow over its surface, with the syncytiotrophoblast having a role in mechanosensing. However, data describing the mechano-sensitive nature of this cell, particularly in early gestation, is lacking. In this study we show that the syncytiotrophoblast expresses six proteins that have been related to shear sensing, and this expression is higher in the first trimester than at term. This suggests shear on the sycytiotrophoblast as an important factor influencing placental morphogenesis early in pregnancy. We then predict shear stress felt by the syncytiotrophoblast in first trimester and term placental tissue using a combination of porous medium modelling and explicit simulations of blood flow in realistic geometries derived from microCT imaging. Our models predict that typical shear stress on first-trimester tissue is higher than at term, supporting the feasibility of this mechanical stimulus as an important driver of healthy placental development.

## 1 Introduction

The placenta is a vital fetal organ that facilitates exchange of nutrients, gases, and wastes between mother and fetus. Pathologies such as fetal growth restriction (FGR), where a fetus fails to achieve its genetic growth potential, are often associated with impaired placental development and function^1^. However, whilst only clinically detectable in later gestation, the pathophysiology of FGR is established in the first half of pregnancy, making this disorder difficult to predict or treat^1^.

In pregnancy, maternal blood flows from the uterine spiral arteries and percolates over the surface of the complex branching villous tree structure of the placenta (in the intervillous space, IVS (Figure 1)). Whilst limited during very early pregnancy, this blood flow increases substantially (15 fold) from the end of the first trimester to term^2,3^. The branching placental villi are covered in a large multinucleated cell, the syncytiotrophoblast, whilst the core of the villi contain feto-placental blood vessels (Figure 1). The feto-placental vasculature grows throughout gestation, and by the third trimester fetal capillaries press against the syncytiotrophoblast to create a thin, efficient, exchange barrier between the physically separate maternal and fetal circulations. Both the volume and velocity of maternal blood that reaches the surface of the placenta, and the development and function of the placenta itself are important determinants of placental exchange capacity and healthy fetal growth.

**Figure 1:**
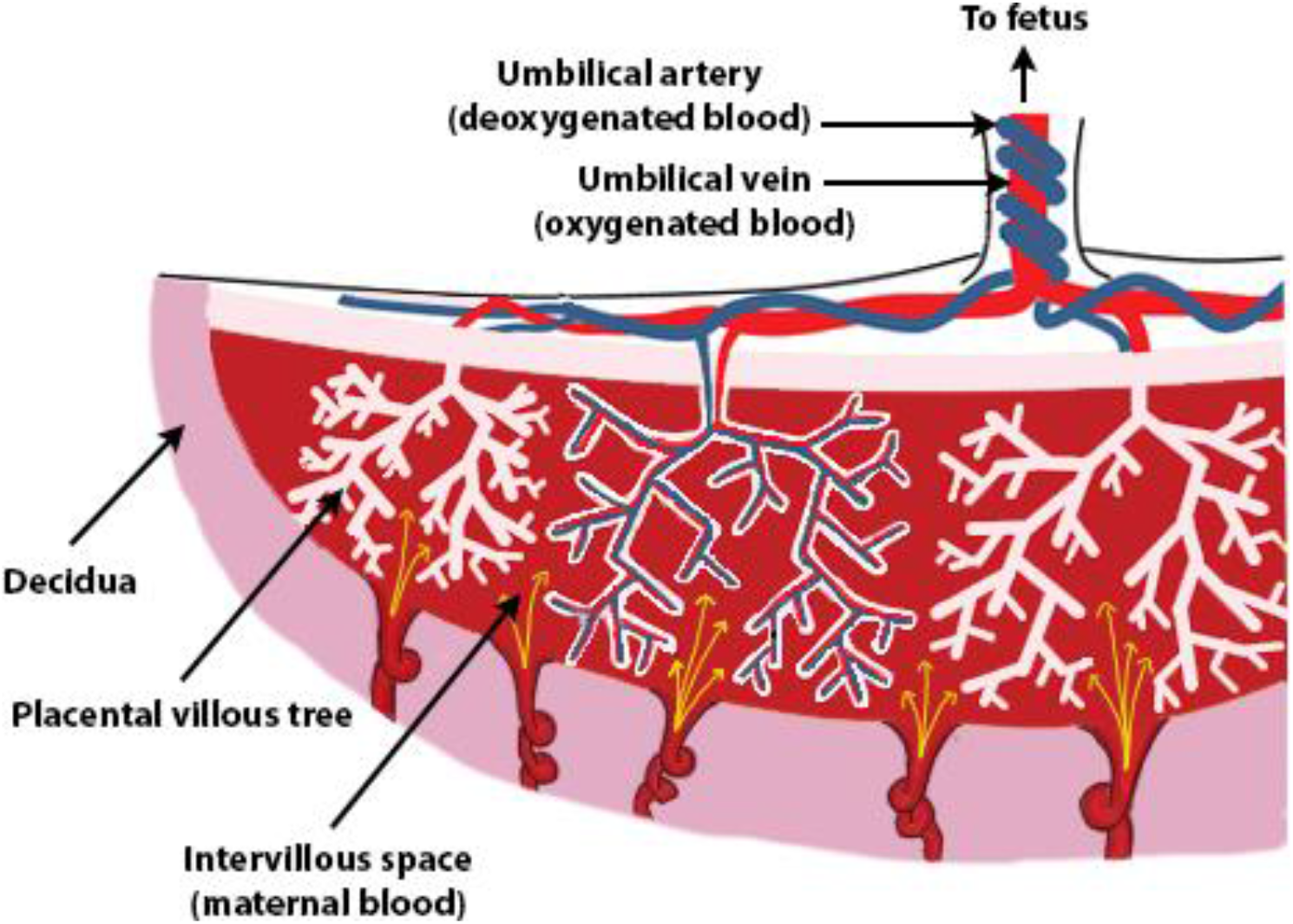
Schematic diagram illustrating the anatomy of the placenta and the two separate circulations, maternal and fetal, that are brought into proximity to each other. The fetal circulation flows within fetal blood vessels that are located within the villous tree, whilst maternal blood circulates around the outside of these trees within the intervillous space.

Recent evidence suggests a relationship between IVS blood flow, syncytiotrophoblast function, and placental development. Using computational modelling and Doppler ultrasound imaging data, Saghian et al.^4^ showed that the penetration of jets of maternal blood into the IVS measured by ultrasound is only possible if ‘central cavities’ of more sparse villi exist near the spiral artery openings^4^. Such central cavities have previously been described anatomically^5,6^, and *in silico* models of placental oxygenation have demonstrated that they act as ‘highways’ ensuring adequate IVS perfusion and oxygen exchange^7^. That central cavities develop progressively throughout gestation, indicates a dynamic relationship between IVS haemodynamics and placental development, with high velocity flow from the spiral arteries impacting regional tissue architecture, and in turn tissue architecture impacting the haemodynamics of local blood flow^4^. As the interface with maternal blood flow, the syncytiotrophoblast is thought to mediate this relationship, and it’s capacity for mechanosensation has been demonstrated both by reports of mechanosensitive protein expression (Polycystin-2 and TRPV6) at term^8,9^, and observations that cultured syncytiotrophoblast cell lines exhibit differential responses in placental growth factor expression and microvilli formation when exposed to fluid shear stress^10,11^. However, there is a need to better understand how the syncytiotrophoblast responds to shear stress, and what shear stresses it is exposed to across gestation.

Mathematical models provide a useful tool to estimate shear stress in the IVS that cannot be measured directly^12,13^. To date these models have focused on term placenta, and have modelled only small volumes of placental tissue^12^, smoothed porous media models^12^, or required time consuming reconstructions of villous tissue architecture from histology^13^. Here, we aimed to extend understanding of shear stress on the syncytiotrophoblast both at term, and at the end of the first trimester (when quantitative analysis of placental structure is more limited). We consider both how the syncytiotrophoblast senses shear stress, and the structure-function relationships that impact the shear stress levels experienced by the syncytiotrophoblast, with an aim to provide insight into how shear stress may impact the dynamic development of the placental villous tree across gestation.

## 2 Methods

First, the expression of mechanosensing proteins by the syncytiotrophoblast was investigated. Then, 3D micro-computed tomography (microCT) images of the surface of placental villous tissue biopsies from first trimester and term placentae were used to quantify tissue architecture. Finally, flow was simulated in the IVS at the placentone (a functional tissue unit fed by a single spiral artery) and villous scales.

### 2.1 Placental Tissue

First trimester placentae (10-13 weeks of gestation) were collected from Auckland Medical Aid Centre. Normal term placentae (38-40 weeks of gestation) were collected from Auckland City Hospital following vaginal or caesarian delivery. The collection of all placental tissue was undertaken following informed consent and approved by the Northern X Ethics Committee (NTS/12/06/057/AM09).

### 2.2 Immunohistochemistry

1cm^3^ blocks of tissue were dissected from first trimester (n=3) and term (n=3) placentae and snap frozen in Optimal Cutting Temperature Compound (OCT) (Agar Scientific, USA). 5μm sections were cut on a cryostat (Leica) and thawed onto poly-lysine coated slides. Sections were fixed in ice cold acetone for 10 minutes, dried, dipped in water, dried, and stored at −20°C until use.

At the time of staining, sections were defrosted, and a boundary marked around the section using a hydrophobic pen (Sigma-Aldrich, USA), then washed with Phosphate Buffered Saline (PBS). 100μL of Blocking Solution (Dako, USA) was added for 10 minutes at room temperature. Wells/slides were washed with PBS-Tween (0.25% Tween in PBS). 100μL of primary antibodies (Table 1) diluted in Blocking Solution were added, and slides were incubated overnight at 4°C.

**Table 1:** Antibodies used for immunohistochemistry

| Antigen antibodies are reactive with | Working Concentration (µg/ml) | Product code and manufacturer |
| --- | --- | --- |
| Dynein-2 | 10 | ab-23905, Abcam, UK |
| IFT88 | 5 | PA5-44087, Thermo-Fisher, USA |
| Kinesin-2 | 8 | ab-15486, Abcam, UK |
| Piezo1 | 20 | PA5-77617, Thermo-Fisher, USA |
| Polycystin-2 | 8 | BS-2158R, Sapphire Bioscience, Australia |
| TRPV6 | 4 | PA5-51295, Thermo-Fisher, USA |
| Irrelevant IgG | 10 | 02-6100, Thermo-Fisher, USA |

The next morning, slides were washed thrice with PBS-Tween. 100μL of biotinylated broad-spectrum secondary antibody solution (Dako, Invitrogen, USA) was added for 1 hour at room temperature. Slides were washed thrice with PBS-Tween, then 100μL of 25μg/mL Streptavidin-488 (Thermo-Fisher, USA) diluted in Blocking Solution was added for 1 hour at room temperature. Slides were washed thrice with PBS-Tween. Finally, nuclei were stained by the addition of 100μL of 10μg/mL Hoechst 33342 (Thermo-Fisher, USA) for 5 minutes at room temperature. Slides were washed with PBS-Tween, and coverslips mounted with Citifluor Antifade mountant (Agar Scientific, USA). Staining was imaged on a Nikon upright fluorescent microscope using NIS software (version 3.2.2). Imaging parameters/microscope settings were held constant between negative control and primary antibodies.

### 2.3 Placental geometry characterisation

#### 2.3.1 Tissue preparation and image acquisition

Explants of first trimester villous tissue (∼1cm^3^) were manually dissected. For term placenta a 0.8cm diameter Bio-Punch (Tedpella, USA) was used to extract a 1.2cm-1.6cm long tissue punch. Tissue was washed gently in PBS then fixed in 4% paraformaldehyde (PFA) in PBS overnight at room temperature. Tissue was then washed in 70% ethanol for 24 hours on a rocker, and subsequently stained with 0.3% phosphotungstic acid (PTA) in 70% ethanol on a rocker at room temperature for 3 days (first trimester tissue) or 9 days (term tissue) to ensure stain permeation to the centre of the sample. Tissue was again washed in 70% ethanol for 24 hours, then embedded in 2% agarose (Sigma-Aldrich, USA) in PBS overnight at 37°C to ensure thermal and chemical stability during imaging.

For imaging, tissue was mounted in a 12mm diameter plastic straw in 2% agarose in PBS, wrapped in 4 layers of Mylar tape, and mounted in the chamber of a Bruker Skyscan 1272 MicroCT. Camera exposure time was tissue-dependent, but within the range of 1500-2000ms. The voltage (kV) and current (µA) for microCT was also sample dependent, but ranged from 75kV:128µA to 85kV:113µA. Samples were rotated 180° during imaging, at a degree step of 0.125°. After the imaging parameters were set, the x-ray beam was turned on, and the sample was kept in the imaging chamber for 45 minutes to allow any initial interaction between the x-rays and sample to subside prior to imaging. No filters except mylar tape were used. Samples were imaged at 3µm/pixel resolution. Imaged volume (excluding tissue edges that may be subject to dissection artefacts) measured approximately 7×7×4mm. Images were reconstructed using NRecon software (version 1.7.4.2, Bruker).

#### 2.3.2 Computational mesh

The process for converting microCT imaging data to a computational mesh is shown in Figure 2, with full details of the segmentation process described in Supplementary Data A. 500×500×500 pixel cubes (1.5×1.5×1.5mm) were extracted from the raw microCT data and segmented into binary images (villous tissue or free space) via a semi-automated procedure.

**Figure 2:**
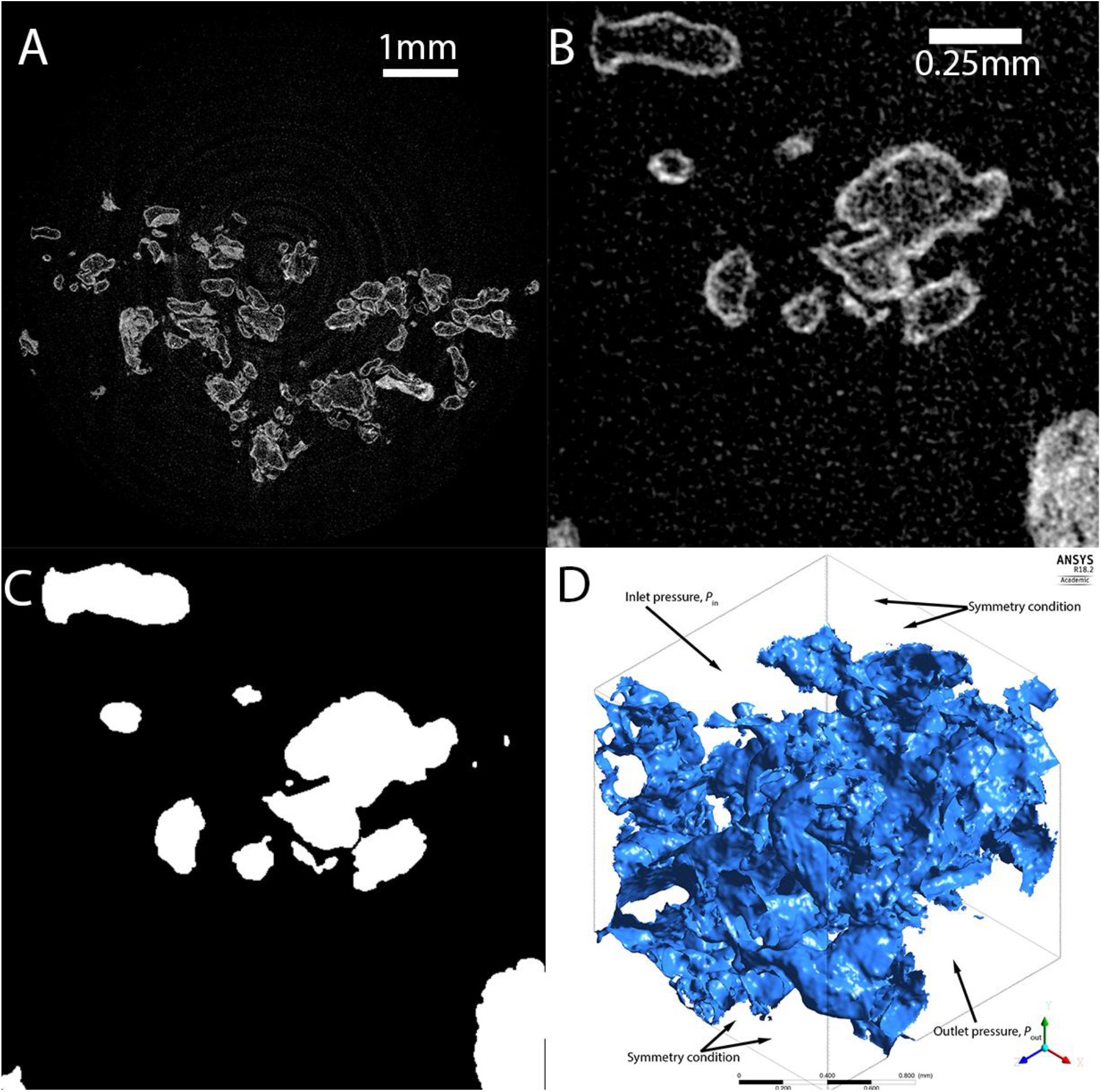
The process of conversion of microCT image data to a computational mesh, using first trimester placenta as an example. (A) MicroCT data covers a volume of approximately 7 × 7 × 4 mm, (B) a 500×500×500 pixel region is extracted from microCT, and (C) villous tissue segmented from this region of interest. (D) Segmented tissue is converted to a computational mesh structure, with boundary conditions covering a 1.5×1.5×1.5 mm block of tissue. In simulation of flow within the tissue blocks pressure boundary conditions are applied to two faces of the cube of tissue, and symmetry conditions are applied at all other faces.

To create a computational mesh, the segmented images were reduced to one quarter size using Matlab (imresize, The Mathworks Inc., 2022a) and then converted into three-dimensional surface meshes using the marching cubes algorithm in the scikit-image python package^14^.

### 2.4 Mathematical model

#### 2.4.1 Placentone (porous media) model

As the blocks of tissue imaged using microCT were not complete representations of the placentone, to simulate these blocks of tissue explicitly, a porous medium model of the whole placentone was employed to determine typical pressure drops over the spatial scale represented by segmented microCT images. A semicircular cap was used to model the placentone with height *Z*_*p*_ and width *X*_*p*_ (Figure 3). Following Saghian et al.^4^, the characteristic funnel shaped opening of the inlet spiral artery was included, with a linear increase in diameter from *d*_*A*_ to *d*_*C*_ over a length of *L*_*B*_. Blood drains through two decidual veins located at a distance 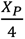 on either side of the spiral artery, with diameter *d*_*V*_. Immediately distal to the spiral artery inlet, there is a villus-free central cavity, taken to be elliptical with height *L*_*CC*_ and width _*d*__*C*_ + 2*d*_extra_.

**Figure 3:**
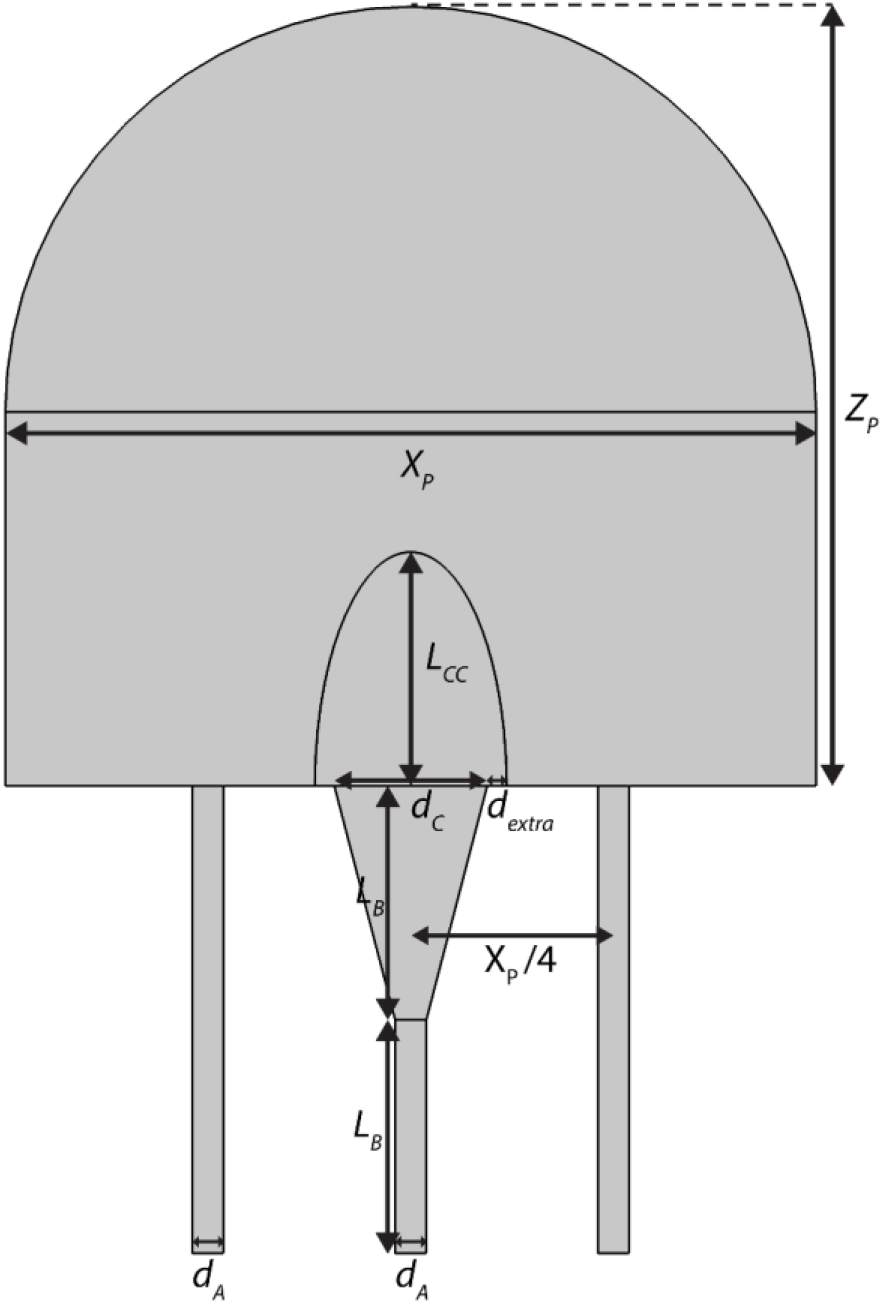
A diagram illustrating the geometry of the placentone model that is used to determine the pressure drop for the explicit flow simulations.

The governing equations for fluid flow are the Navier-Stokes equations which are applied in the central cavity, the inlet spiral artery, and outlet decidual veins. Everywhere else, the Darcy-Brinkman and continuity equations were applied, representing flow of a viscous fluid in a porous medium (Supplementary Data B). The placentone model was parameterised to both term and first trimester characteristics based on Saghian et al.^4^ (Table 2). One parameter, *ϕ*, the porosity of the IVS is available from microCT data, and a second parameter, *k*, its permeability is derived from porosity by the Kozeny–Carman equations^4,15^. Villus diameter is required for permeability calculation; at term we used 70μm^4^ and in first trimester 200µm ^16^. The range of porosities measured from our microCT images was 0.37-0.52 (term) and 0.71-0.85 (first trimester). These values compare to a literature range of 0.3–0.7 at term^15,17^, and whilst there are no direct reports in the first trimester, estimates suggest a porosity of up to 0.92^4^. An inflow velocity condition at the spiral artery boundary was applied, based on ultrasound, as described by Saghian et al.^4^.

**Table 2:**
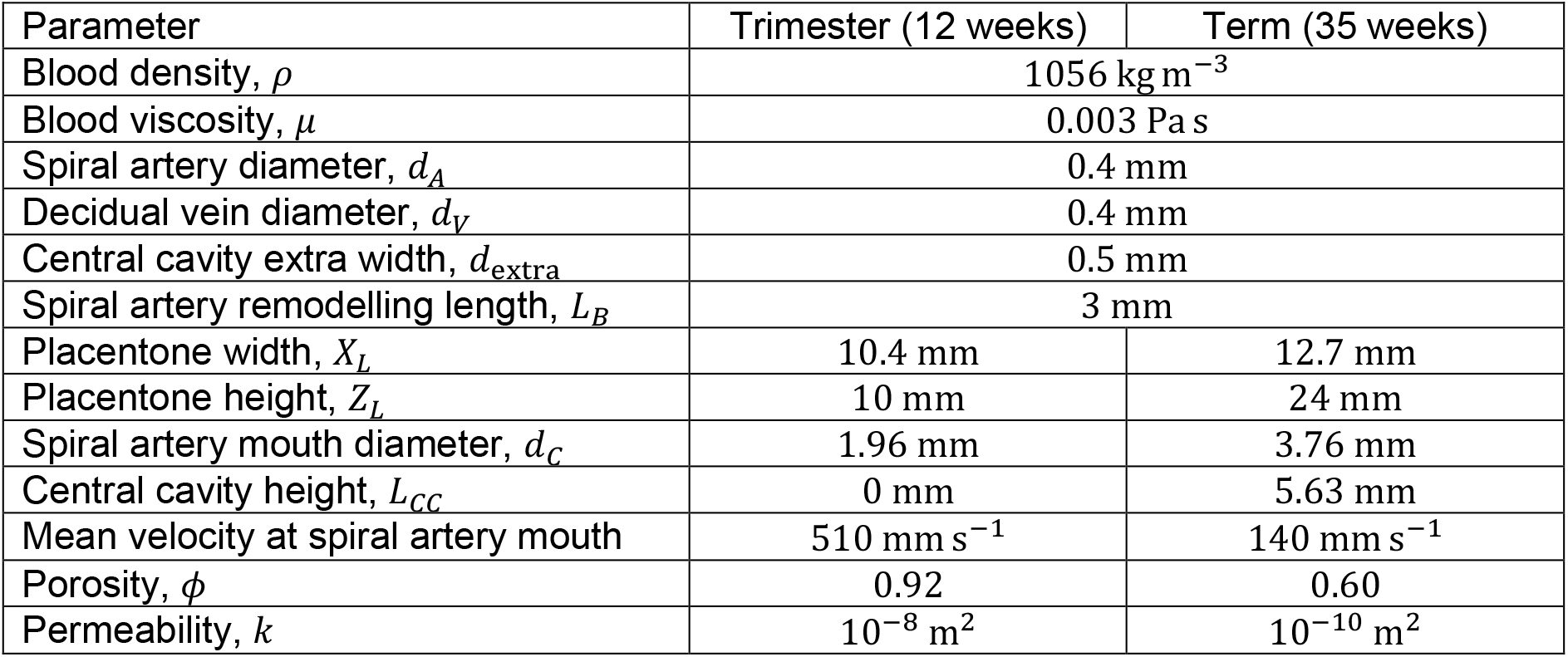
Parameter values used for the first trimester or term porous media models.

| Parameter | Trimester (12 weeks) | Term (35 weeks) |
| --- | --- | --- |
| Blood density, $\rho$ | 1056 kg m<sup>-3</sup> | |
| Blood viscosity, $\mu$ | 0.003 Pa s | |
| Spiral artery diameter, $d_A$ | 0.4 mm | |
| Decidual vein diameter, $d_V$ | 0.4 mm | |
| Central cavity extra width, $d_{\text{extra}}$ | 0.5 mm | |
| Spiral artery remodelling length, $L_B$ | 3 mm | |
| Placentone width, $X_L$ | 10.4 mm | 12.7 mm |
| Placentone height, $Z_L$ | 10 mm | 24 mm |
| Spiral artery mouth diameter, $d_C$ | 1.96 mm | 3.76 mm |
| Central cavity height, $L_{CC}$ | 0 mm | 5.63 mm |
| Mean velocity at spiral artery mouth | 510 mm s<sup>-1</sup> | 140 mm s<sup>-1</sup> |
| Porosity, $\phi$ | 0.92 | 0.60 |
| Permeability, $k$ | 10<sup>-8</sup> m<sup>2</sup> | 10<sup>-10</sup> m<sup>2</sup> |

#### 2.4.2 Villous tree model

To model the shear stress exerted on villous tree, flow was explicitly simulated in the IVS in geometries derived from microCT using the Navier-Stokes equations (Supplementary Data B). On the domain boundaries, one set of opposite faces was taken to be the inlet and outlet faces, and symmetry conditions were applied to the remaining four faces (see Figure 2D)^18^. As the inlet and outlet faces do not have a uniform boundary (due to the presence of the villous structures), a uniform pressure at the inlet and outlet was prescribed, rather than velocity. Hence the flow within the IVS is driven by the pressure drop across the volume of placental tissue. The Reynold’s number determined for the problem implied that the Navier-Stokes equations could be simplified to linear Stoke’s equations (explored numerically in Section 3.3). As the pressure drop across a volume of placental tissue is difficult to measure experimentally,and would vary in magnitude across the placenta, the placentone scale porous media model was used to determine this pressure drop.

### 2.5 Computational methods

#### 2.5.1 Porous media model

The porous media model was implemented in COMSOL Multiphysics (COMSOL Inc., 5.2a) using the laminar flow solver with porous media domains. The geometry was meshed in COMSOL and convergence was determined by mesh refinement. The mean velocity at the spiral artery mouth was computed using the line average integration function. To determine the distribution of pressure gradients within the placentone, the pressure gradient was sampled over a uniform grid and linear interpolation. The central cavity (in the term placenta), spiral arteries, and veins were excluded from sampling. Sampled pressure gradients were used to create a normalised histogram of pressure gradients (a probability density). The number of sample points was increased until a converged histogram was achieved.

#### 2.5.2 Villous tree model

Villous tree surface meshes were imported into Ansys ICEM (Ansys Inc., 2021 R2) and unstructured tetrahedral meshes created. Simulations were carried out using Ansys Fluent (Ansys Inc., 2021 R2) with convergence determined by monitoring the convergence of the wall shear stress experienced by the villous tree, as well as the maximum vertex velocity present on the inlet surface, and stopping the simulation when the relative change between iterations was less than 10^−3^. Mesh convergence was determined by a mesh refinement study.

#### 2.5.3 Combined model

Due to the linearity of the explicit flow simulations (Stokes flow), the pressure gradient distribution from the porous media model was combined with the distribution for shear stress for a given pressure gradient. The centre of each bin in the pressure gradient histogram was used to scale the shear stress histogram from explicit simulations. The resulting histogram could then be normalized by the probability density of the pressure gradient bin in question.

Repeating this process for each bin and adding the results created a new combined histogram for shear stress distribution that accounts for the modelled distribution of pressure gradient in the placentone.

## 3 Results

### 3.1 Syncytiotrophoblast expression of mechanosensing proteins is higher in first trimester than term

To more comprehensively profile mechanosensing protein expression in the syncytiotrophoblast, and how this changes across gestation, the expression of six different mechanosensing proteins was determined by immunofluorescence. Dynein 2, IFT88, and Kinesin 2 are microtubular motor-proteins that facilitate the biogenesis and/or maintenance of primary cilium^19,20^, whilst Piezo1, Polycystin-2, and TRPV6 are mechanosensitive ion channels that facilitate Ca2+ influx into the cell in response to shear stress^21-23^. All stained uniformly and strongly in first trimester syncytiotrophoblast (Figure 4, n=3 for each). However, at term syncytiotrophoblast staining for all six proteins was weaker, limited to discrete regions of the syncytiotrophoblast, and more heterogenous, showing varied expression both within and between villi in the same placenta (Figure 4). Regardless of gestation, staining for Dynein-2, IFT88 and Kinesin-1 was limited to the syncytiotrophoblast, with no staining observed in cytotrophoblast, mesenchymal cells, or placental endothelial cells (Figure 4). In contrast, staining for Piezo1, Polycystin-2 and TRPV6 was also observed in placental endothelial cells (Figure 4). No staining was observed in sections incubated with irrelevant IgG antibodies (negative controls).

**Figure 4:**
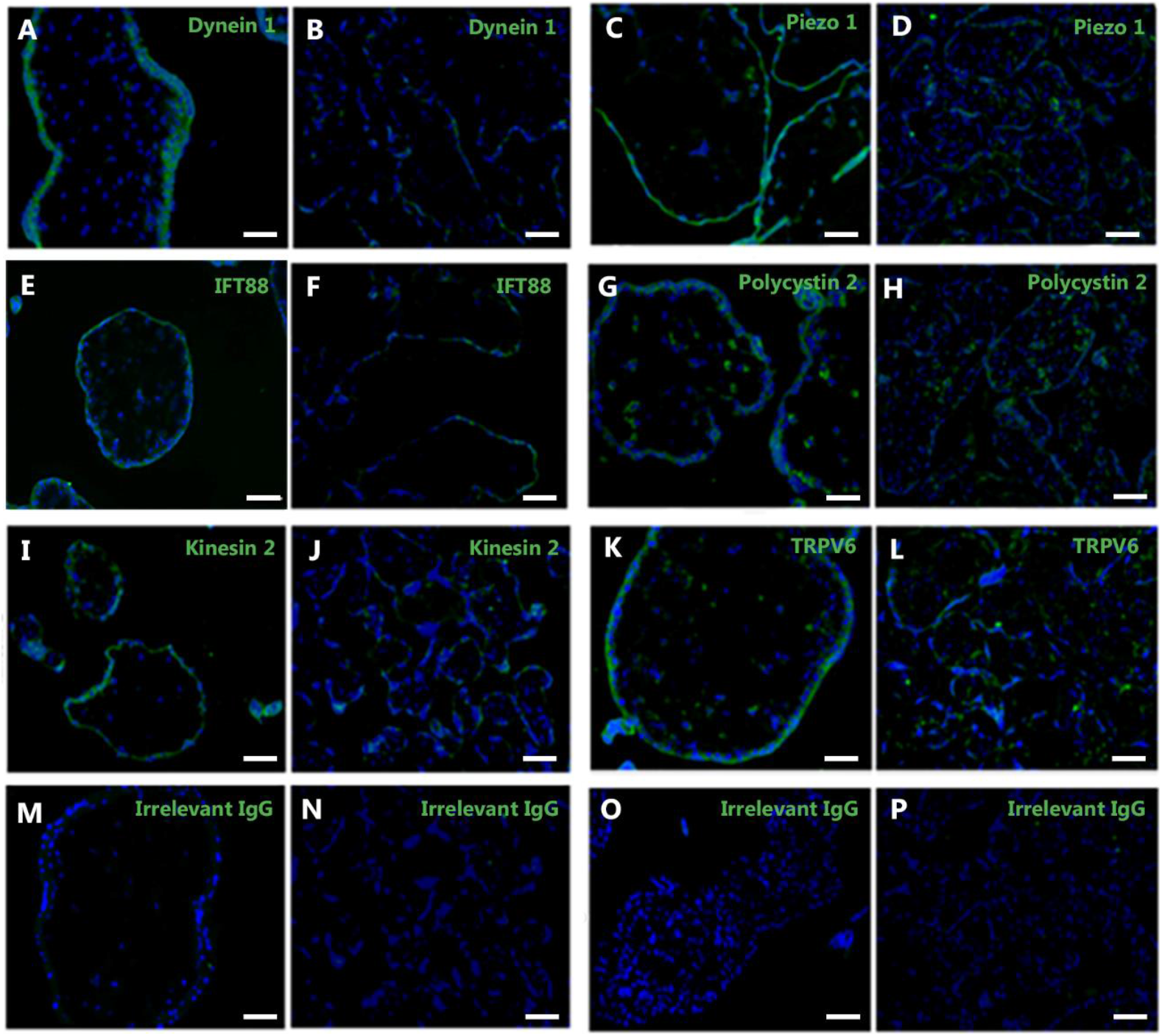
Photomicrographs of immunofluorescent staining of late first trimester (A, C, E, G, I, K, M, O) or term (B, D, F, H, J, L, N, P) placental tissue with antibodies reactive with the motor proteins Dynein 1 (A, B), IFT88 (E, F), or Kinesin 2 (I,K), or the mechanosensitive ion channels Piezo 1 (C, D), Polycystin-2 (G, H), or TRPV6 (K, L). Sections stained with irrelevant IgG antibodies were used as negative controls (M-P). Nuclei are counterstained with Hoescht 33342 (blue). Scale bars = 100µm.

### 3.2 Prediction of driving pressure throughout the IVS using a placentone level model

To predict the range of pressure gradients in the first trimester and at term, we used a placentone scale porous media model (Figure 5). Comparing the velocities (Figure 5A,B), we observe the jet penetration length expected from ultrasound data^24^, and previously predicted computationally^4^. The term placentone experiences high pressure gradients near the decidual veins, and this is higher than in the same region in the first trimester (Figure 5C,D). However, overall, the term placenta has a distribution of pressure gradients that are lower than the first trimester scenario. This is because of the larger size of the term placentone, meaning that more villous tissue is located far from decidual veins (thus experiencing small pressure gradients).

**Figure 5:**
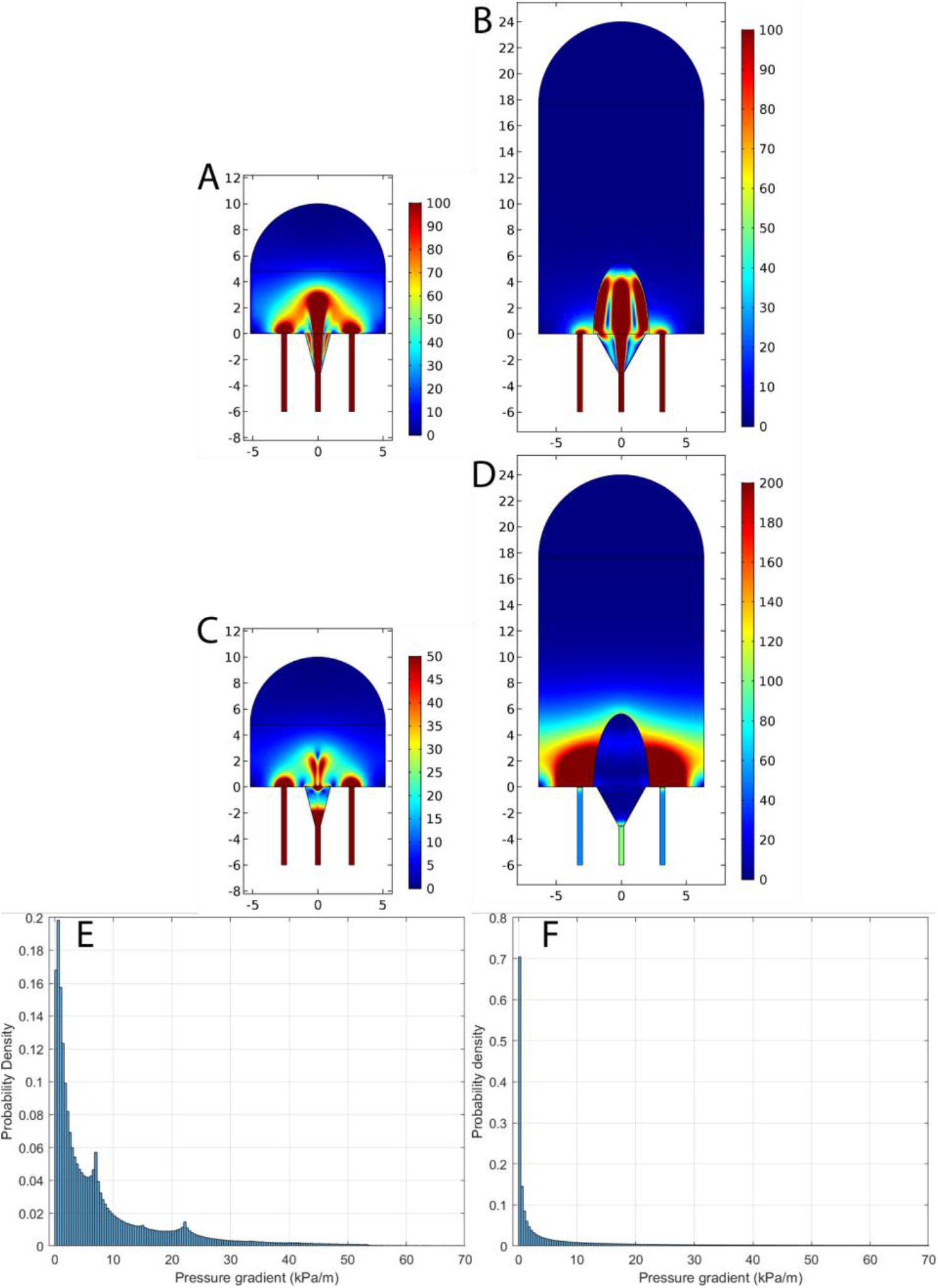
Velocity (A,B), pressure gradient magnitude (C,D), or distribution of pressure gradient magnitude (E, F) for flow simulated around a first term placentone unit (A,C,E) or a term placentone unit (B, D, F). Note that colour scales in (C,D) are not the same.

### 3.3 Prediction of shear stress experienced by the syncytiotrophoblast across gestation

To determine the shear stress experienced by the syncytiotrophoblast at the villous scale we simulated flow through the microCT-imaged IVS of first trimester and term placenta using pressure gradients predicted by the placentone model. Local flow and shear stress was predicted with a pressure gradient of 1 kPa/m (1.5 Pa pressure drop) applied in the x-direction, (Figure 6). All simulations were then repeated with a pressure gradient applied in the y- and z-directions, and although this leads to some variation in the shear stress distribution, this variation is much smaller than the variation between individual samples at the same stage of pregnancy, or across gestation (Supplementary Data C).

**Figure 6:**
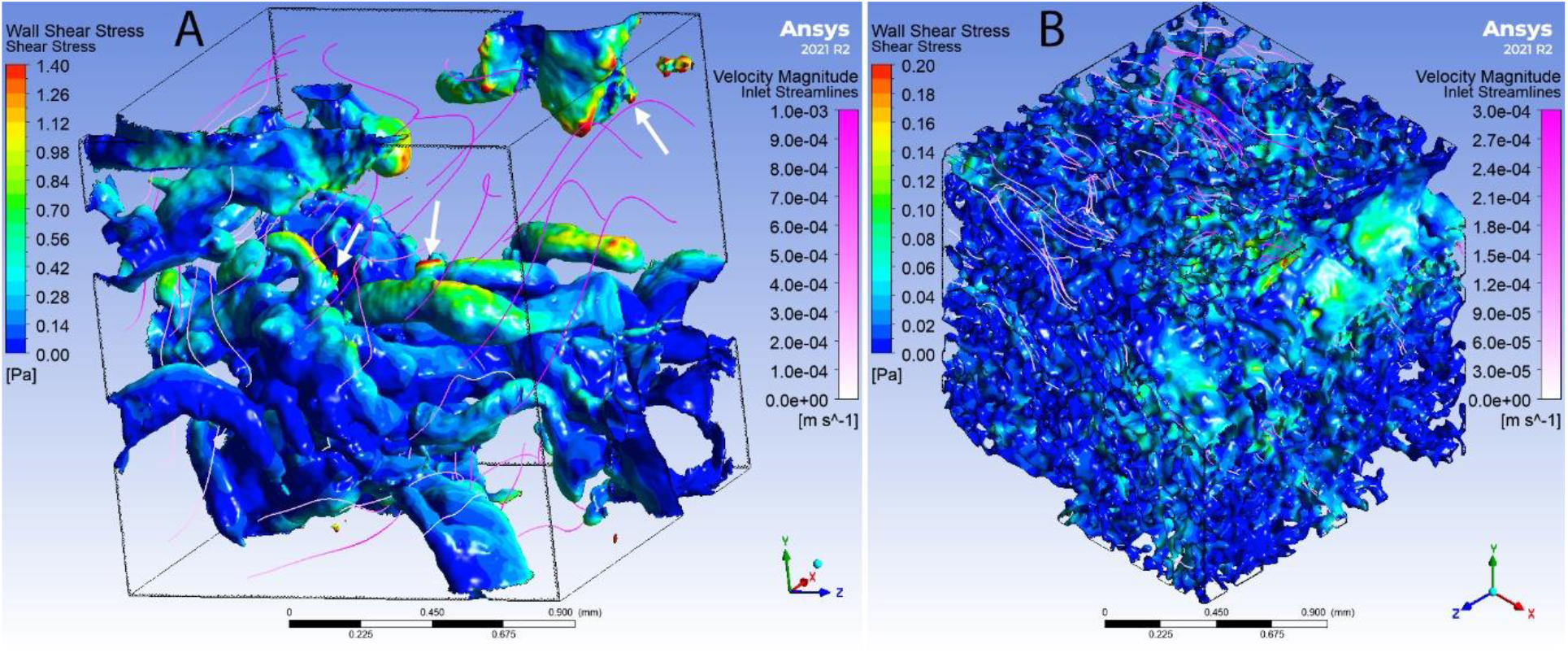
Examples of model predictions of shear stress from the explicit simulations for (A) first trimester or (B) term placental tissue, including wall shear stress and fluid flow streamlines. Results are for a pressure gradient of 1 kPa m−1 which corresponds to a pressure drop of 1.5 Pa over our 1.5 mm sample.

Boxplots of shear stress distribution for each sample with a pressure gradient of 1 kPa/m are shown in Figure 7. For the same pressure gradient, the first trimester samples are predicted to experience a higher shear stress than term samples. The shear stress experienced is dependent on the specific geometry of the placental segments assessed (comparing. samples of the same type, Figure 7). Combining the data, first trimester tissue is expected to experience a median shear stress of approximately 0.026 Pa compared to 0.0054 Pa for term tissue when subjected to a pressure gradient of 1 kPa/m.

**Figure 7:**
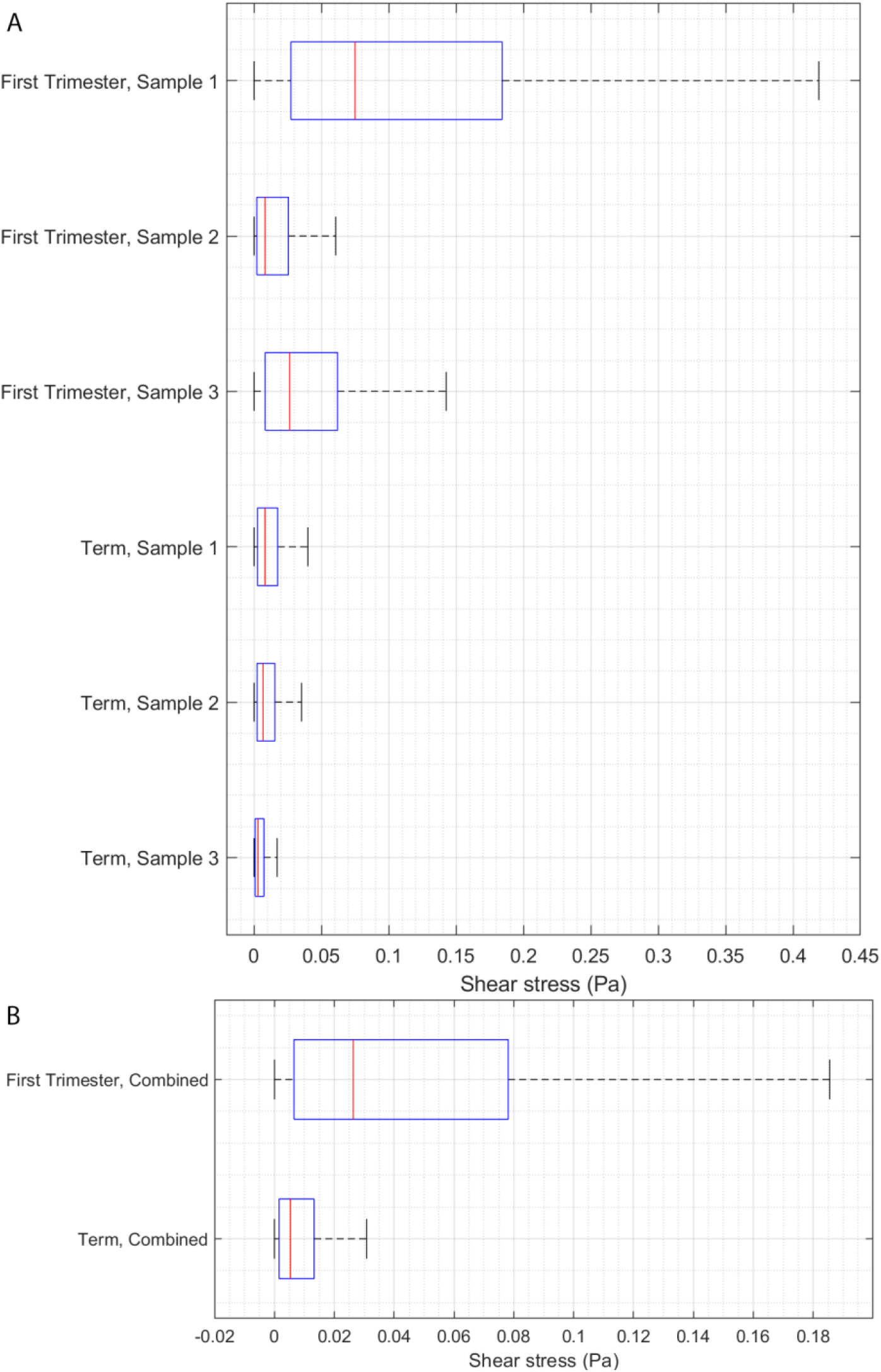
Boxplots of shear stress obtained from the villous tree model simulations for (A) the individual samples of first trimester and term placenta, (B) combined distribution of all first trimester samples and all term samples. All results correspond to a pressure gradient of 1 kPa/m.

### 3.4 Accuracy of the Stokes approximation

We expect low Reynold’s number throughout most of the IVS, which allows for Stokes flow approximations. Stokes flow allows scaling of simulation results, meaning only a single simulation needs to be run to obtain shear stress distributions under a variety of pressure drops. To gauge the accuracy of this approximation, we performed a numerical study on one sample of term placenta (Supplementary Data D). We find that the Stokes approximation holds with input pressure gradients of up to 400 kPa/m, within the expected range seen in Figure 5 (C,D,E,F).

### 3.5 Combined model

Having obtained a distribution of pressure gradients from the placentone model and the distribution of shear stress for a given pressure gradient from the villous tree model, these two distributions were combined to obtain an overall distribution of shear stress for each type of placentone. Figure 8 shows the overall predicted distribution of shear stress for first trimester and term as well as their cumulative densities. Similar to our villous level simulations in 3.3, a higher level of shear stress is predicted for first trimester tissue compared to term. Cumulative density plots suggest the first trimester placentone experiences a higher median shear stress (0.08 Pa (0.8 dyn/cm^2^)) compared to term (0.004 Pa (0.04 dyn/cm^2^)). Furthermore, 90% of the placentone experiences shear stress less than 1.3 Pa (13 dyn/cm^2^) for first trimester compared to 0.25 Pa (2.5 dyn/cm^2^) for term.

**Figure 8:**
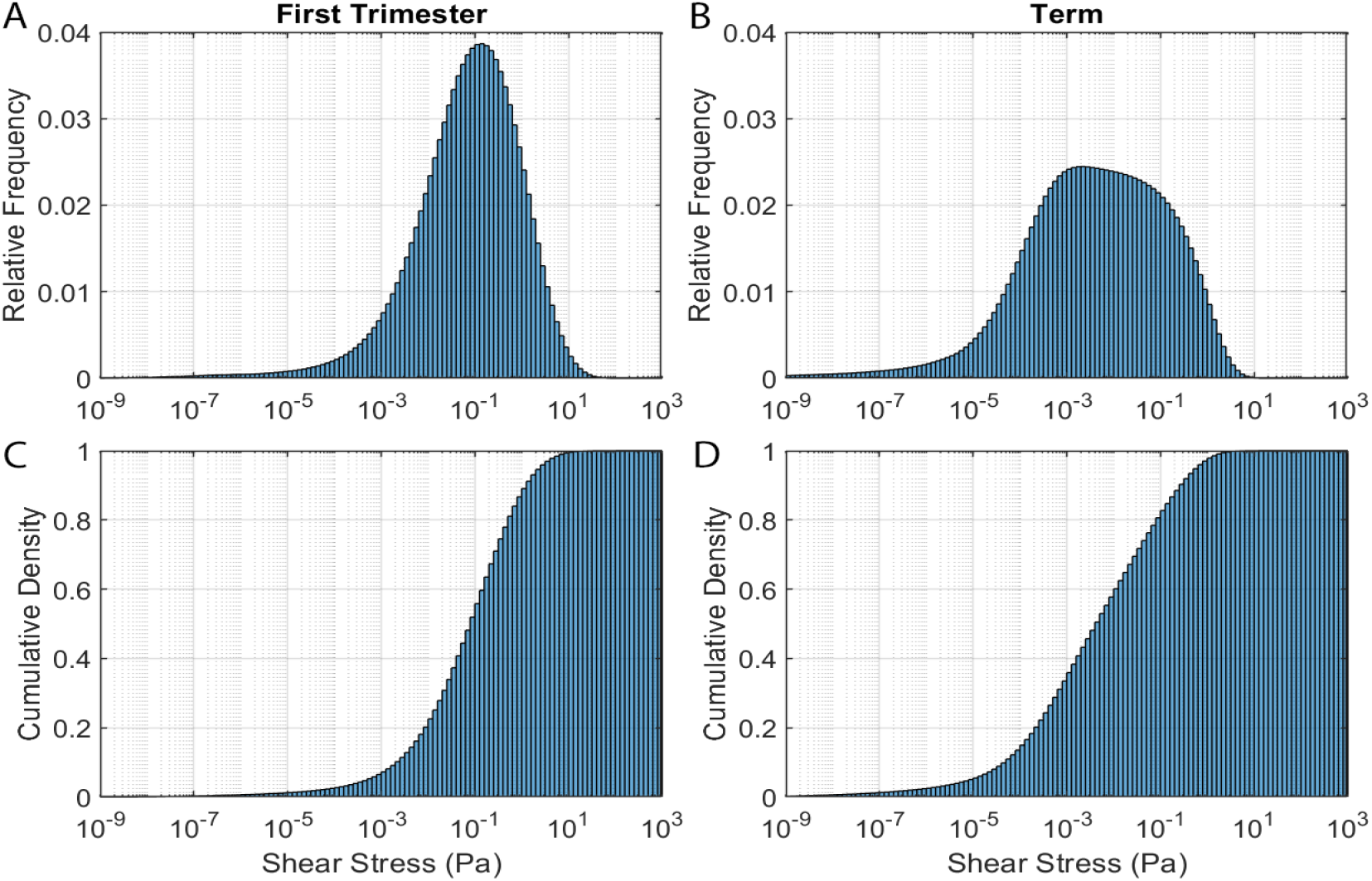
A combined histogram showing the (A, B) relative frequency distribution, (C, D) cumulative density that takes into account both the distribution of pressure gradient within the placenta and the distribution of shear stress for a given pressure gradient for (A,C) first trimester and (B,D) term placenta.

## 4 Discussion

In this study, we sought to better understand the impact of shear stress on the surface of the placenta across gestation by demonstrating how the syncytiotrophoblast may sense shear stress, and predicting the shear stress it is exposed to. Higher expression of mechanosensing proteins in the syncytiotrophoblast of late first trimester placentae (compared to term) demonstrated that IVS flow may influence early placental morphogenesis. This highlighted the need to understand the range of shear stresses on the syncytiotrophoblast at the end of the first trimester, and computational models were then used to predict this for the first time. Together, these models provide new understanding of how structure-function relationships between villous architecture and IVS blood flow may impact syncytiotrophoblast shear stress across gestation.

The syncytiotrophoblast expressed all six mechanosensitive proteins examined, demonstrating capability for mechanosensation throughout pregnancy. Whilst polycystin-2 expression has previously been demonstrated at term^8^, and TRPV6 expression has been demonstrated on syncytialised BeWos and in membrane fractions from term syncytiotrophoblast^9,11^, this data significantly increases our scope of understanding of the range of both motor proteins and mechanosensitive ion channels expressed by the syncytium. In particular, syncytiotrophoblast-specific expression of motor proteins that are normally associated with primary cilia in other tissues (Dynein-2, IFT88, Kinesin-2)^19,20,25^ suggests that, in addition to their role increasing surface area for nutrient/gas transport, microvilli on the placental surface may also be involved in mechanosensation.

Mechanosensitive protein expression has not previously been examined in first trimester placental tissue. Here, all mechanosensing proteins examined were expressed at a higher level in the first trimester than term. This highlights the important impact that the onset of maternal blood flow through the IVS may have on placental development and function from the end of the first trimester, and supports our previous data suggesting a relationship between IVS blood flow and placental villous development from the end of the first trimester^4^. Mechanosensation impacts tissue morphogenesis in a range of other tissues^26^. Recently, TRPV6 was shown to be important for extracellular matrix formation in the murine placental labyrinth^27^, and in morphogenesis and tissue folding in pig placentae^28^, highlighting its relevance to placental tissue remodelling. Finally, as the syncytiotrophoblast expresses nitric oxide synthase (NOS)^29^, our data expanding the scope of expression of mechanosensitive ion channels that induce nitric oxide (NO) production suggests that the syncytiotrophoblast may produce NO in response to shear stress, like endothelial cells do^21,22^. Syncytiotrophoblast NO production has been hypothesized to play a role in decreasing feto-placental vascular resistance at the onset of IVS blood flow at end of the first trimester^30^, and in preventing platelet activation and adherence to the syncytiotrophoblast surface, which can lead to fibrin deposition and impair exchange^31^.

To truly understand the impact of shear stress on placental function, it is important to determine the normal range of shear stresses the syncytiotrophoblast is exposed to *in vivo*, and mathematical modelling provides a vital tool to do this. Lecarpentier et al.^12^ followed the same placentone scale approach employed in this study, and others^4,15^, and also conducted explicit 2D simulations of flow in geometries created from a single histological section of term placenta (providing macro-scale estimates of shear stress), and over single terminal villous imaged in 3D using scanning electron microscopy (providing micro-scale information on peaks in local shear stress). However, the two models are disconnected and did not simultaneously consider variation across scales. Roth et al.^13^ used time consuming reconstructions of serial histological sections of a smaller 2mm region of villous tissue immediately distal to a spiral artery from a single FGR placenta, which was perturbed to assess the impact of spiral artery haemodynamics. Here, we combined a porous medium model at the placentone scale with explicit simulations in geometries derived from microCT (providing 3D tissue architecture). Tun et al.^32^ used synchrotron X-ray imaging in 8 mm^3^ tissue blocks stained with zinc-based fixative Z7 and PTA to simulate flow in the IVS in small, cropped regions, but considered a much smaller region of tissue (0.2 mm^3^) than in this study. Synchrotron imaging is high-resolution compared with conventional microCT, but far less accessible due to cost and size requirements of facilities. The micro-CT imaging employed here provides an efficient means to analyse multiple samples of tissue, at relatively large volumes, without excessive computational expense (our simulations ranged from 70-140 hours on 16-cores (Intel(R) Core(TM) i7-7700 CPU at 3.60 GHz).

Lecarpentier et al.^12^ predicted a gradient in shear stress on the syncytiotrophoblast at term as one moves distally from the spiral artery mouth into the IVS. Their data suggested a range in mean shear stress over a single terminal villous between 0.05-0.23Pa (based on a local maternal blood velocity range of 0.1-1.0mm^3^/s). We estimate that 90% of villi within a placentone at term experience a shear stress <0.25Pa, which is of similar magnitude to the upper estimate in that study. Our estimate for median shear stress (0.004Pa) is below their expected range, which may relate to our placentone model having a lower permeability at term. We estimated permeability based on villous architecture, and Lecarpentier et al. calculated permeability following an analysis that aimed to match flow velocities derived from magnetic resonance imaging. Roth et al.^13^ did not directly report shear stress ranges, although their data visualisations suggest a shear stress of <5Pa in the normal term placenta. As their model geometry reflects only the 2mm of tissue closest to the spiral artery, this is comparable to our highest predicted shear stress near the uterine blood vessels feeding the IVS, and our model suggests the same order of magnitude predictions in that region.

No previous study has predicted first trimester shear stress on the syncytiotrophoblast. We predict that first trimester tissue experiences a higher level of shear stress than term tissue for the same driving pressure. This is likely due to a sparser tissue architecture in the first trimester, providing a lower resistance to flow. Driving pressure predicted in the placentone model allowed determination of shear stress at the villous scale. Our model predicts the highest driving pressure near model outlets (decidual veins), with highest peak driving pressures at this location in term placentae. However, overall, the term placentones larger size means that it experiences lower pressure gradients. This translates to a lower median value of shear stress at term in our combined model, despite localised regions of higher shear near to the spiral artery mouth. The 90th percentile values that we predict (1.3Pa (13dyn/cm^2^) for first trimester and 0.25Pa (2.5dyn/cm^2^) for term) illustrate that parts of the villous trees may experience a much higher shear stress compared to the median value. Future work relating this spatial variation in shear stress across the IVS to syncytiotrophoblast mechanosensation/transduction is required to elucidate what impact this may have on villous morphogenesis and placental function.

Local villus level shear stress information can also be derived from our computational method. In our first trimester images, we noted protrusions from villi (Figure 6A, white arrows) that are possibly either syncytial sprouts (newly forming villi), or syncytial nuclear aggregates (multinuclear clusters of aged syncytiotrophoblast shed throughout gestation). Our computational results predict these locations as sites of elevated shear stress. As such, it is interesting to speculate on the role of elevated shear stress in promoting villous branching from existing ‘hot spots’ (regions of existing villi with high cell proliferation^33^), or in driving syncytiotrophoblast turnover.

Our model assumes a Stokes approximation in explicit simulations that allow a small number of simulations to be scaled to produce predictions of shear stress distribution within the whole placentone. By analysing the Stokes approximation, we find it begins to break down only in regions very close to the outlet veins in the first trimester model, and in a slightly larger portion near the veins in the term model. However, these locations make up a negligible portion of the overall distribution of pressure gradient at the whole placentone level, and so in general the Stokes assumption holds. These regions, however, would potentially contribute to localised high shear stress, and our model thus may underestimate absolute maximum shear stress.

In conclusion, by combining a porous media model of a placentone with explicit simulations from samples of villous tissue, our computational model allows us to predict the overall shear stress distribution experienced by the placenta across gestation at multiple, linked scales. We show that the first trimester syncytiotrophoblast both exhibits an increased expression of mechanosensitive proteins, and a higher average shear stress than the term placenta, suggesting an important role for shear stress in placental development from early gestation.

## Acknowledgements

This work was supported by Auckland Bioengineering Institute Faculty Research Development fund and a Health Research Council of New Zealand Consolidator Grant (20/926).

