## Supplementary Material for "Multi-scale modelling of shear stress on the syncytiotrophoblast: Could maternal blood flow impact placental function across gestation?"

### Supplementary Data A: Image segmentation

Subsamples of 500x500x500 pixel (corresponding to cube sections of 1.5 mm edge length) sections were extracted from the samples for segmentation. Due to differences in the density of tissue and intensity of staining between the first trimester and term samples, the image segmentation process was slightly different in each case. However, the process was similar, and a semi-automated segmentation was possible in each case. Segmentation was conducted using ImageJ and the python SimpleITK package.

#### *First trimester segmentation*

Images were despeckled by application of a 3x3x3 pixel median filter. The image was then thresholded using to produce a binary image, with the threshold manually selected by the user to distinguish between background (intervillous space) and tissue. There were gaps in the walls in the binary thresholded image, and some unfilled interiors due to lack of staining in the core of relatively large villi (phosphotungstic acid staining primarily highlights the villous surface and the fetal blood vessels within the villous tree, not the mesenchymal stroma). Gaps in the walls were corrected manually, slice by slice. Any small holes in the interior of the tissue were then filled a binary hole filling algorithm and the images were cleaned of random noise by removing all connected components with size less than 5 pixels.

#### *Term segmentation*

Images were despeckled by application of a 3x3x3 pixel median filter. Then, an alternating sequential filter which is a sequence of opening and closing using binary spheres of radius 1 and 2, which aims to smooth the image while maintaining as much as possible the villous surface topology. As with the first trimester gaps in the walls in the binary thresholded image, and some unfilled interiors due to lack of staining in the villous core. In these images, both gaps in the wall and unstained villous tissue were manually corrected, slice by slice. Finally, small holes in the interior of the tissue were then filled a binary hole filling algorithm and the images were cleaned of random noise by removing all connected components with size less than 5 pixels. The major difference between image analysis in the term tissue compared to the first trimester tissue was in the requirement of an additional alternating sequential filter to suppress noise and additional manual correction, particularly in large villous structures where staining of the mesenchymal stroma was not sufficient to be captured by automated segmentation algorithms.

### Supplementary Data B: Governing equations

The governing equations for flow in the cotyledon (except in the central cavity, the inlet spiral artery, and outlet decidual veins) were taken to be the Darcy-Brinkman and continuity equations

$$\frac{\boldsymbol{\rho}}{\boldsymbol{\phi}^2}(\boldsymbol{u}\cdot\nabla\boldsymbol{u}) = -\nabla\boldsymbol{P} + \frac{\boldsymbol{\mu}}{\boldsymbol{\phi}}\nabla^2\boldsymbol{u} - \frac{\boldsymbol{\mu}}{\boldsymbol{k}}\boldsymbol{u}, \tag{B.1}$$

$$\nabla\cdot\boldsymbol{u} = \boldsymbol{0}. \tag{B.2}$$

where $u$ and $P$ represent the fluid velocity and pressure respectively and $\rho$ and $\mu$ are the density and dynamic viscosity of blood in the intervillous space respectively, $\boldsymbol{\phi}$ is the porosity of the intervillous space and $\boldsymbol{k}$ is its permeability. In the central cavity, spiral artery, and decidual veins, blood is free to flow unhindered and hence the fluid flow in these regions is governed by the Navier-Stokes equations, (B.3).

To model the shear stress exerted on villous tree, we explicitly simulate flow in the intervillous space in geometries derived from microCT. The surface of the villous tree that is treated as an impermeable boundary. Assuming that blood in the intervillous space behaves as a Newtonian fluid, the governing equations for fluid flow in the intervillous space are given by the Navier-Stokes and continuity equations (under steady conditions)

$$\rho \cdot (\boldsymbol{u} \cdot \boldsymbol{\nabla u}) = -\boldsymbol{\nabla} P + \mu \nabla^2 \boldsymbol{u}, \tag{B.3}$$

$$\boldsymbol{\nabla} \cdot \boldsymbol{u} = 0, \tag{B.4}$$

The steady-state approximation has been assessed in previous models [20], and found to be well founded. A no-slip boundary condition is applied on the tissue surface

$$\boldsymbol{u} = 0. \tag{B.5}$$

The Reynold's number was considered as to inform whether a full Navier-Stokes treatment of the flow was needed. Taking the characteristic length to be on the order of magnitude of the intervillous spacing (pore size), $\boldsymbol{L} = \mathbf{10^2}\ \mu\mathbf{m}$ together with $\boldsymbol{u} = \mathbf{10\ mm\,s^{-1}}$, $\boldsymbol{\rho} = \mathbf{10^3\ kg\,m^{-3}}$ and $\boldsymbol{\mu} = \mathbf{1\ mPa\,s}$ gave a Reynolds number of $\boldsymbol{Re} = \frac{\rho u L}{\mu} = \mathbf{0.1}$, suggesting that for typical physiological values of flow through the intervillous space, we are in the Stokes regime in which we can neglect the inertial term, and hence equations are linear. This will remain true so long as the velocity remains small enough. However, as the pressure drop across the domain is increased, this velocity will increase, potentially violating the Stokes approximation.

### Supplementary Data C: Impact of Boundary conditions in explicit simulations

To evaluate the effect of the direction of the pressure boundary conditions on the results from the villous tree model, we performed additional simulations with the pressure drop applied in the y and z-directions for Sample 1 of each type (pressure drop was applied in the x-direction in the original simulations of all samples). Results from these simulations can be seen in Figure S1. Based on these results, we find that although changing the direction of the applied pressure drop does alter the predicted shear stress, this variation is much smaller than the variation that occurs between samples and between tissue types.

### Supplementary Material D: Validity of Stokes flow assumptions

To test the validity of the assumption that our solution is linear with respect to the applied pressure gradient as it is well approximated by Stokes flow (due to the low Reynolds number), we performed additional simulations for term tissue (Sample 1) with different applied pressure drops. Results from these simulations can be seen in Figure S2. These results show that this assumption of linearity holds up to at least 400 kPa/m and we only begin to see a small divergence from this trend at 4000 kPa/m.

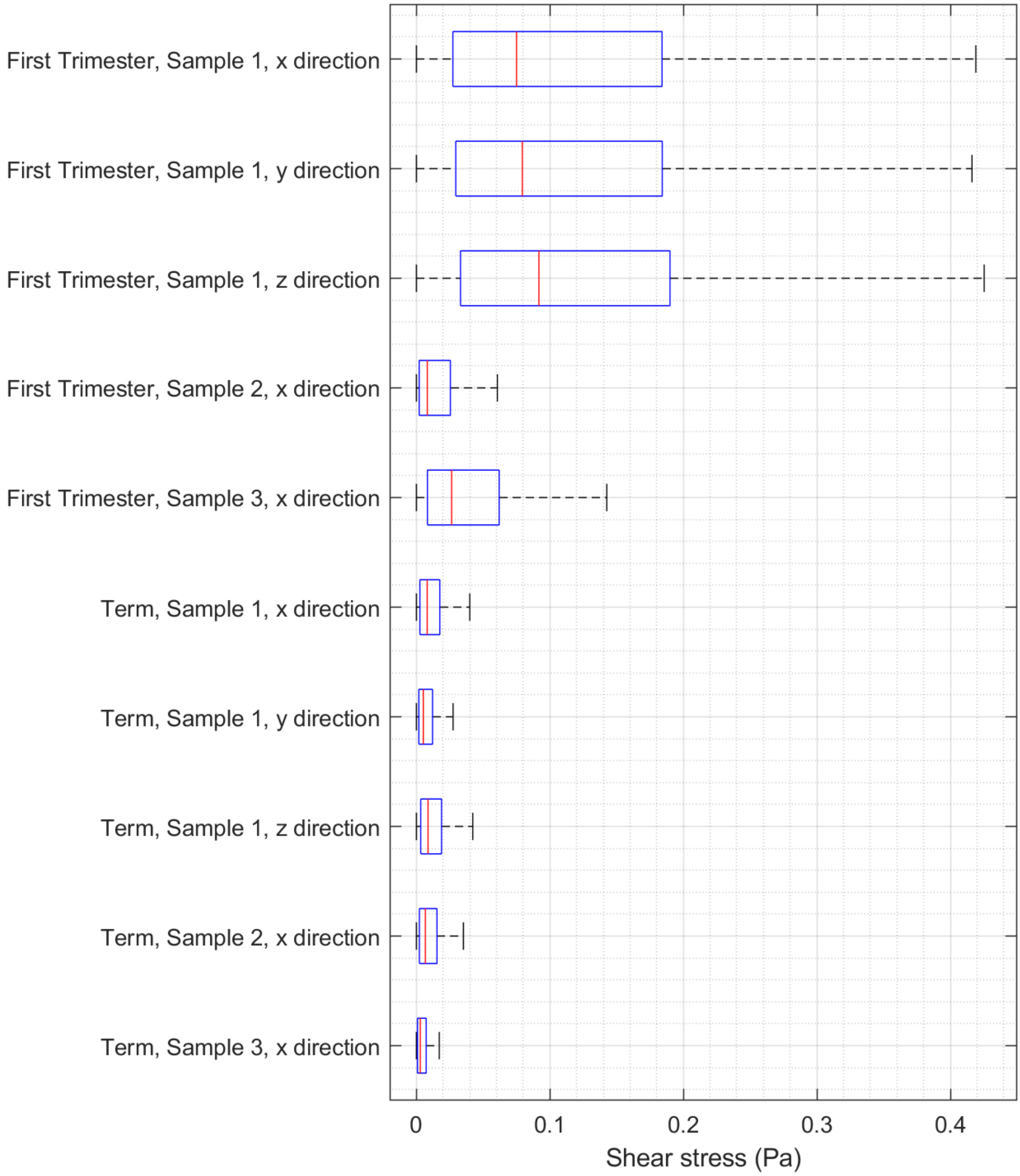


*Figure S1: A boxplots of shear stress obtained from the explicit fluid simulations with additional simulations for Sample 1 of each type that evaluate the effect of the direction of the applied pressure boundary conditions. All results correspond to a pressure gradient of 1 kPa/m. Note: Results for the x-direction of all samples are the same as the results given in Figure 7 and are repeated here to allow easy comparison.*

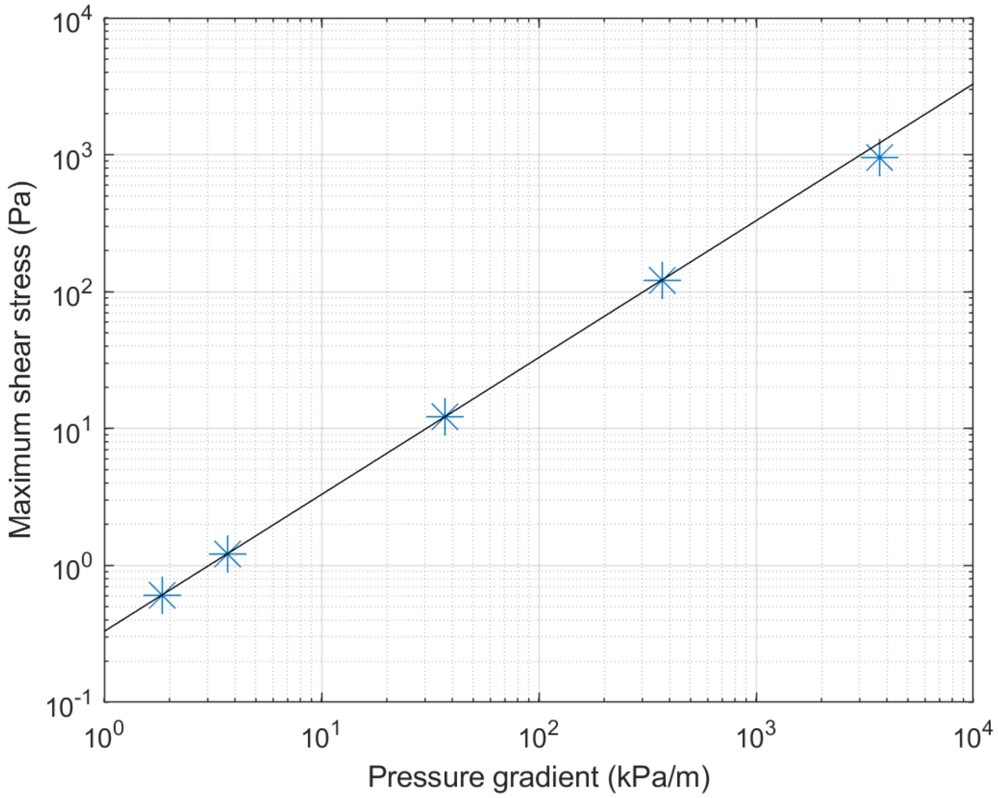


*Figure S2: A plot of predicted maximum shear stress for a term placenta with flow in the x direction against the input pressure gradient. Simulation results are shown as crosses while the black line represents the expected shear stress given the linearity of the solution that comes from the Stokes approximation. We see that the Stokes approximation holds up to at least a pressure gradient of about 400 kPa/m.*
